# Enhancing the Representational Similarity Between Execution and Imagination of Movement Using Network-Based Brain Computer Interfacing

**DOI:** 10.1101/166603

**Authors:** Neda Kordjazi, Amineh Koravand, Heidi Sveistrup

**Author notes:** Neda Kordjazi was a Graduate research associate at the University of Ottawa at the time of this study.

## Abstract

Motor imagery-based brain computer interfacing (MI-BCI) as a neuro-rehabilitation tool aims at facilitating motor improvement using mental practice. However, the effectiveness of MI-BCI in producing clinically meaningful functional outcome is debated. Aside from computational shortcomings, a main limiting obstacle seems to be the substantial representational dissimilarity between movement imagination (MI) and movement execution (ME) on the level of engaged neural networks. This dissimilarity renders inducing functionally effective and long lasting changes in motor behavior through MI challenging. Moreover, the quality and intensity of imagination is highly prone to change on a trial-to-trial basis, based on the subject's state of mind and mental fatigue. This leads to an inconsistent profile of neural activity throughout training, limiting learning in a Hebbian sense. To address these issues, we propose a neuroconnectivity-based paradigm, as a systematic priming technique to be utilized pre-BCI training. In the proposed paradigm, ME-idle representational dissimilarity network (RDN) features are used to detect MI in real-time. This means that to drive the virtual environment, an ME-like activation pattern has to be learned and generated in the brain through MI. This contrasts with conventional BCIs which consider a successful MI, one that results in higher than a threshold change in the power of sensorimotor rhythms. Our results show that four out of five participants achieved a consistent session-to-session enhancement in their net MI-ME network-level similarity (mean change rate of 6.16% ± 4.64 per session). We suggest that the proposed paradigm, if utilized as a priming technique pre-BCI training, can potentially enhance the neural and functional effectiveness. This can be achieved through 1- shifting MI towards engaging ME-related networks to a higher extent, and 2- inducing consistency in MI quality by using the ME-related networks as the ground-truth and thus enhancing the robustness of the activity pattern in the brain. This would in turn lend to the clinical acceptability of BCI as a neurorehabilitation tool.

## 1. Introduction

The goal of movement imagination-based brain-computer interfacing (MI-BCI) as a neurorehabilitation tool is to use mental practice as a proxy to guide neuro-plastic reorganization in the lesioned brain towards relearning motor skills [1, 2]. This idea is based on the premise of shared neural correlates of movement imagination (MI) and overt movement execution (ME) [3–6]. Following the execution of the movement, sensorimotor feedback is provided to the brain through afferent tracts [7]. The afferent feedback is an essential component of human motor control, both in error-based and reinforcement aspects of learning [8, 9]. This feedback is missing in MI, which is defined as the state of actively planning a movement, without execution. MI-BCI provides a pathway to close the sensorimotor loop by providing feedback to the user whenever an MI is detected [10, 11]. Such a technique would be most beneficial to individuals with severe disabilities who demonstrate little benefit from physical exercise [12–14]. However, despite extensive research, MI-BCI for motor neurorehabilitation is far from becoming a standardized clinical paradigm.

In this study we will address two main obstacles limiting the clinical applicability of BCI. First, from a network point of view, different connectivity patterns have been associated to MI versus ME, casting doubt on the effectiveness of MI being a suitable replacement for ME in motor rehabilitation. To be more specific, motor planning/imagination of both left and right hand movements mainly engages the left posterior parietal and left motor areas [15–18]. However, manual actions are accompanied by a bilateral inter-hemispheric connection between the motor areas [19, 20], while the descending tracts originate from the primary motor cortex contralateral to the active hand [21]. This network level representational dissimilarity hinders the practicality of using MI as a proxy to induce motor-related plasticity in the brain.

Most MI-BCI systems currently used for motor neurorehabilitation rely on features derived from individual EEG channels, especially the changes in the sensorimotor rhythms in terms of spectral band power (BP) [17, 22, 23]. Whenever this change exceeds a certain threshold, a successful MI is registered and subsequently the subject receives a feedback. Although BP features offer minimal computational workload as well as fair accuracy for real-time BCI, they inherently overlook the task-related interplay of neuronal activity in the engaged neural networks. However, the short and long-term changes in the engaged networks as a result of training, can be considered a proxy to assess the neuroplasticity induction process [24, 25]. With such a measure in hand, one can monitor the process of learning, and possibly modify training appropriately on a session to session basis [26–28].

Moreover, unlike active movement which is easily controllable, the quality as well as intensity of MI can vary on a trial-to-trial basis [29, 30]. Active movement training thus results in a consistent and robust exercise paradigm, with predictable outcome in certain muscle groups. MI however, is an endogenous mental task. Although one can be instructed to imagine movement in a certain way, e.g. kinesthetically, the exact way in which the instructions translate into action is highly subjective and difficult to measure [31].

Movement imagination thus results in an inconsistent and dispersed profile of neural activity, throughout the training period, which negatively affects the net functional benefits of training. Moreover, the immense variability in MI might partly explain the large heterogeneity in the ability of both healthy and clinical populations in controlling α-or β-range BP MI-BCIs [32, 33]. This calls for a much needed “ground-truth map” to be used as the gold-standard reference of comparison for MI validation. The most logical choice for such a reference would be the brain activity during ME.

To address the above-mentioned issues, a fundamental revision of the BP-based MI detection frameworks typically used in MI-BCI is critical. One theoretically suitable substitute for BP seem to be the measures of neural network connectivity. The idea mainly originates from the fact that the brain is undoubtedly an interconnected network, with distinct patterns of inter-regional information flow for different tasks (see the review by [34]). Neural network connectivity can be looked at on two main levels; anatomical (structural) level and statistical (functional and effective) level [35]. The changes in the structural level usually happen on a slow time scale (hours or days) and therefore are not suitable for BCI whereas statistical connectivity changes on a millisecond timescale [36, 37].

### 1.1. Network-based BCI

Functional and effective connectivity are two main data-driven approaches to look into statistical dependencies in multichannel recordings. Functional connectivity provides information on the correlation or mutual dependency between recording channels, whereas effective connectivity looks into the partial predictive power of the activity in one region over other regions [34]. Functional and effective connectivity have both been applied to BCI algorithms. Daly and colleagues [38] aimed at separating single cued executed and imagined finger taps versus resting state to control a BCI system, using 19-channel EEG data and measures of functional connectivity. They used Hidden Markov models to model the temporal dynamics of the mean clustering coefficients at each frequency bin. They achieved a subject mean accuracy of 70.9% for executed and 67.3% for imagined movements, suggesting the reliability of functional correlations in EEG, as a measure for decoding ME and MI.

Liu et al [39] used a nonlinear Granger Causality (GC) measure of functional connectivity and a statistical threshold in the context of a BCI system. Different intended arm reaching movements (left, right and forward) were decoded using 128 EEG surface electrodes. The authors showed that directional information flow patterns were distinct for each intended arm movement direction. This study suggests that statistical connectivity can be used as a differentiating measure for movement intention even when the same end effector is used.

Billinger et. al. [40] applied independent components analysis (ICA) to 45-channel EEG data to separate imagined movement from resting state. They did so before estimating measures of effective connectivity from single-trial vector autoregressive (VAR) models. They reported that effective connectivity measures extracted from independent components can achieve accuracy similar to band-power features, however they did not use their proposed technique in a real-time MI-BCI setting. Additionally, using ICA to eliminate inter-channel dependencies is in contrast with the concept of effective connectivity which looks for cause and effect relations (delayed dependencies) between recordings.

Heger et. al. [41] proposed a feature extraction method for BCI applications by combining connectivity measures as feature level filters for power spectral density (PSD) features, and evaluated the performance of this method in a finger tapping 65-channel EEG dataset and a 118-channel motor imagery dataset. The authors were able to improve classification accuracy compared to spectral band power, confirming the applicability of statistical connectivity measures to real-time BCI.

However, there are two challenging factors that have not been systematically investigated: one computational and one conceptual. On the computational side, connectivity extraction from EEG signals usually requires high spatial density recordings (20 or more channels). Thus the algorithmic workload of a connectivity-based BCI system can be quite high, rendering the system slow. On the other hand, if the number of recordings is insufficient, the system might face lack of information and become unstable. On the conceptual side, reinforcing the neural networks contributing to MI, without taking into account MIME representational differences, will not necessarily help improve the functional outcome of BCI training.

To address both of the above-mentioned challenges, we propose and test a priming technique to be used pre-BCI training, which itself consists of BCI training. We perused two main goals in designing the BCI paradigm. First we wished to ascertain the feasibility of implementing a network-based BCI feature that offers low computational workload, provides robust accuracy, demands low channel density recording and is completely self-paced (not cued), meaning participants are given the full benefit of using their own intent. Second we aimed at investigating the effectiveness of using such BCI training in changing the quality of MI to engage ME-related networks to a higher extent and moreover, increase the MI robustness by enhancing MI-ME representational similarity.

To this end, the connectivity-based paradigm utilizes ME-idle representational dissimilarity network (ME-idle RDN) characteristics as features to detect MI in real-time. This means that to drive the virtual reality interface, an ME-like activation pattern has to be generated through MI in the brain. This is unlike conventional BCI which defines a successful MI, one that results in a change in the BP of the sensorimotor rhythm exceeding a threshold[42]. Incremental levels of difficulty are considered in the design of the training paradigm, i.e. as training progresses, the MI-ME representational similarity network (RSN) has to become stronger to be able to drive the environment. We hypothesize that the proposed paradigm will result in a shift in MI, towards engaging ME-related networks of the same movement to a higher extent.

## 2. Methods

### 2.1. Subjects

Five right-handed healthy subjects were recruited (mean age = 38.4 years, SD = 16.5, range 26–55, 4 female). The participants had no habitual drug or alcohol consumption, cognitive or psychiatric impairments, neurological disorders, metal implants or pregnancy. Subjects were compensated for their participation and gave their written, informed consent beforehand. The study protocol was approved by the University of Ottawa Health Sciences and Science research ethics board and the Bruyère research ethics board.

### 2.2. EEG recording

In all experiments, EEG was recorded from 32 Channels (AFz, AF3, AF4, F3, F1, Fz, F2, F4, FC5, FC3, FC1, FCZ, FC2, FC4, FC6, C5, C3, C1, Cz, C2, C4, C6, CP5, CP3, CP1, CPz, CP2, CP4, CP6, P3, Pz, P4) grounded to Fpz. All measurements were performed at a sampling rate of 1000Hz, using a Brain Products Amplifier, and it was made sure that the electrode-skin impedance was kept below 5KΩ in all electrodes. The data were transmitted online to Matlab via a secure IP connection for real-time analysis. All programs used in this study for online and offline processing were custom coded by the first author either in Python or Matlab.

### 2.3. Assessment of kinesthetic and visual imagery

For the assessment of motor imagery vividness we used the short version of the Kinesthetic and Visual Imagery Questionnaire (KVIQ-10). KVIQ-10 uses a five-point scale to rate the clarity of image (V subscale) and the intensity of sensations (K subscale) during imagination of five small movements of individual limbs. An examiner first explained the differences between kinesthetic and visual imagery, as well as every movement on the list to each subject. Subjects were then required to rate their imagery using the operational definition of each category (e.g., 5 = an “image as clear as seeing” for visual, or “feeling as clear as doing” for kinesthetic). All subjects were instructed to use their non-dominant hand (left hand) for the KVIQ imagery tasks. Subjects scoring 3 or lower in any of the tasks were excluded from the study.

### 2.4. Connectivity analysis

Measures of effective connectivity were used for feature-extraction from the EEG data to drive the BCI as well as offline network analysis. Effective connectivity provides a measure of interaction between remote brain regions using the predictive power of the activity in one region to explain the activity of other regions [43]. Multiple measures of effective connectivity have been introduced and applied to neural data such as Granger causality (GC) and its multiple extensions [39, 44], partial directed coherence (PDC)[45], transfer entropy [46] and directed transfer function (DTF) [47].

Most effective connectivity measures are extracted from the vector autoregressive (VAR) models coefficients. MVAR describe how time series in a multichannel data depend on past values of each other. As such, VAR models relate to the concept of causality, meaning that their structure mirrors the temporal order of driver and effect relations among the time series also referred to as direction of information flow. Assuming the time series is stationary, an MVAR model of order p can be written as shown in (1).

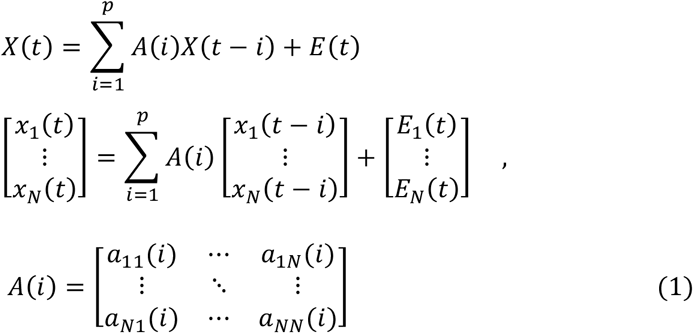

where *X(t)=[X_1_(t) X_2_(t) … X_N_(t)]^T^* represents an N-channel dataset, *A(i)* is the *N×N* matrix of autoregressive coefficients and *E(t)* is the vector of residuals or prediction error. In the frequency domain, (1) can be written as shown in (2) to (4).

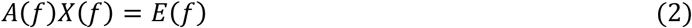

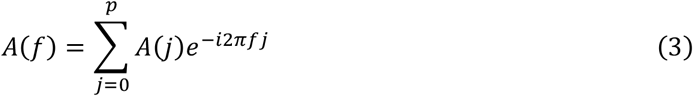

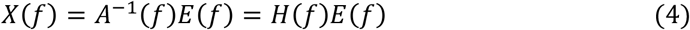

where *A(0) = −I* (unity matrix). Then the cross spectral density is obtained as shown in (5).

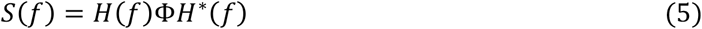

where *Φ* is the covariance matrix of the error vector *E(t)*. Using the above equations we can derive information flow extent and direction, between time series using various measures. In this study DTF was used for network generation because of its robustness with respect to systematic variation of signal to noise ratio, length of the recording data [48], optimal choice of the MVAR model order and sampling frequency [49]. *DTF* is calculated as follows:

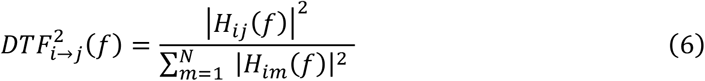

Where 0 ≤ *DTF_i→j_*(*f*) ≤ 1 and 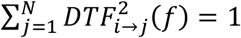 for all *1≤j≤N*. *DTF* measures connectivity by focusing on the net information outflow from channel *i* to channel *j*, normalized by the influence of channel *i* on all other channels.

Given that EEG data is highly non-stationary, the methodology used for autoregressive coefficient estimation, as well as data window size is crucial to the validity of results obtained by measures like *DTF*. In addition, EEG also suffers from multicollinearity [50], i.e. the existence of near-linear relationships among channels mainly because of noise and volume conduction of the head. During regression calculations, this relationship can create inaccurate estimates of the regression coefficients, inflate the standard errors of the regression coefficients, and degrade the predictability of the model.

To address this issue, we first filtered the signal with a spline interpolation spatial Laplacian filter to minimize instantaneous correlations between channels [51]. In addition we employed Ridge Regression (RR) for model parameter estimation, a technique for analyzing multiple regression multidimensional data with multicollinearity [52]. When multicollinearity occurs, least squares estimates are unbiased, but their variances are large so they may be far from the true value. By adding a degree of bias to the regression estimates, ridge regression reduces the standard errors.

### 2.5. Study design

The study consisted of three phases: (1) screening session, (2) subject-dependent number of BCI training sessions and (3) one retention session, 10 days after the last training session (Figure 1).

**Figure 1:**
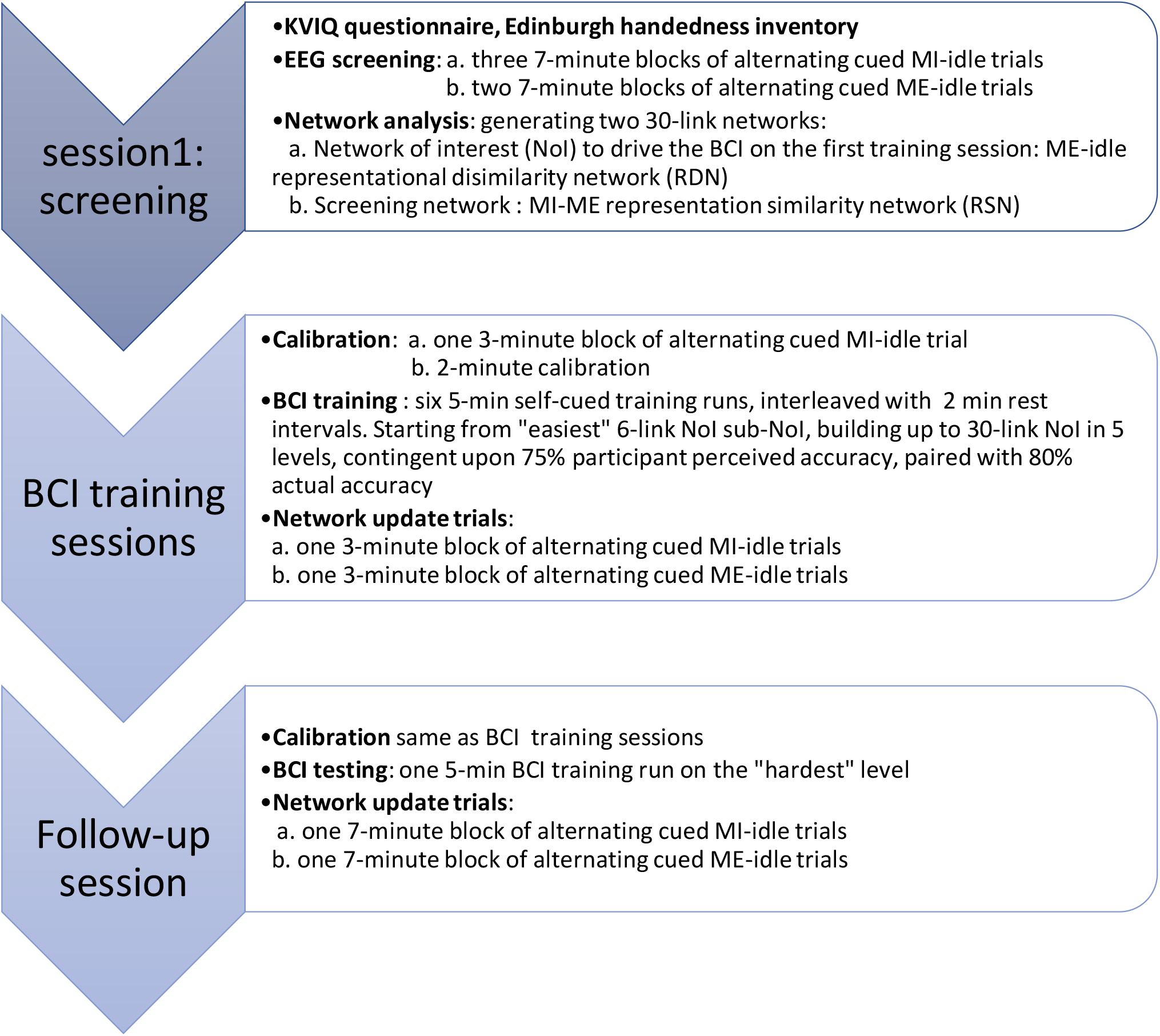
Flow-chart view of the MI-BCI paradigm

#### 2.5.1. Screening session

Subjects were asked to fill out the KVIQ-10, as well as the Edinburgh handedness inventory. EEG screening was then performed with the subject seated comfortably on a chair in a shielded, dimly lit room in front of a display set at a 1-meter distance. A virtual reality hand avatar, developed in-house using VRML was used as the visual cue. The virtual hand opened and closed with the repetition cycle of 2 seconds (1 sec to open and 1 sec to close, Figure 2). An auditory GO/STOP cue was used to cue participants to imagine movement (GO cue) or rest (STOP cue). When the GO cue was played, the avatar hand opened and closed for 8 seconds. The movement stopped with the STOP cue, and the avatar remained in resting position for 8 seconds, until the next GO.

**Figure 2:**
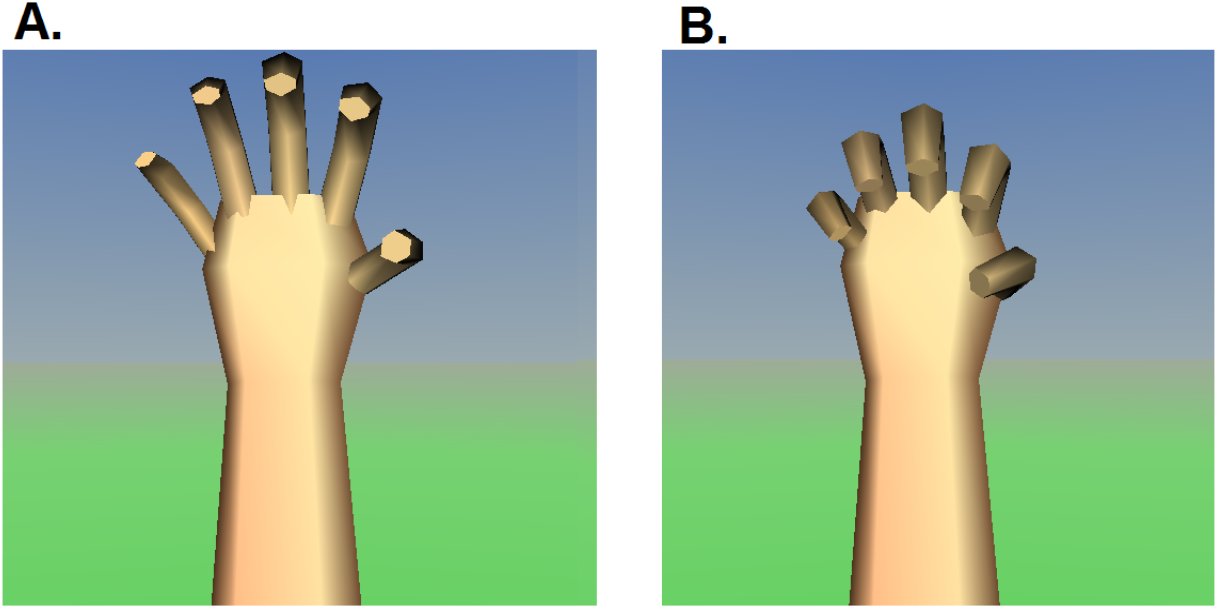
The hand avatar used as the visual cue in fully open (A.), and fully closed (B.) positions

Subjects were asked to look at the visual cue on the display (Figure 2) accompanied with the auditory cue, and imagine the kinesthetic feeling of the same movement as the avatar in their own non-dominant hand. Each block lasted for 7 minutes. Three 7-minute blocks of alternating MI/idle trials, interleaved with 2 to 3-minute resting intervals were recorded with the same protocol. Subjects were then asked to complete two additional 7-minute blocks of alternating ME/idle trials, where they had to actively open and close their hand at the same pace as the avatar movement. The screening session took about 50 minutes.

##### Generating the representational similarity/dissimilarity networks

Screening session EEG data were used to generate two 30-link networks. First, the ME-idle representational dissimilarity network (RDN) was generated and subsequently used to drive the BCI. Hereafter, this network will be called the Network of Interest (*NoI*). The second network generated was the MI-ME representational similarity network (RSN) for screening. To generate these networks, EEG data corresponding to ME, MI and idle states in each 8-sec trial were segmented into single-second long non-overlapping windows (N windows per block for each state). The connectivity matrices were then calculated for each data window (*N* matrices of dimensions 32 × 32 per state). Element *c_ij_* (*i* ≠ *j*) connectivity matrix, represents the partial predictive power of channel *i* over channel *j*.

In order to generate the *NoI*, a generalized linear model (GLM) was fitted to the off-diagonal elements of the connectivity matrices in an R-fold leave-one-block-out cross-validated fashion (R the total number of blocks). Considering the large number of connections and the limited amount of data in hand, within-subject cross-validation is critical to ensure that the model is not over-fit, and has satisfactory generalization to unseen data. To this end, in each fold, all except one block were considered the training set, and the left-out block was considered the testing set. For each block, the mean connectivity pattern, was calculated by averaging each off-diagonal element over the N windows for each of the two states of ME and idle (32^2^ − 32 = 992 off-diagonal elements, *c_ij_* (*i* ≠ *j*)). This resulted in a *992×1* vector of average connectivity pattern per state, per block. The GLM was then fitted as follows:

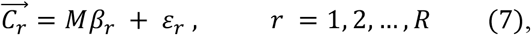

where 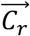 is the vector obtained from concatenating the two 992 × 1 vectors (1984 × 1) of mean connectivity patterns for the two states, averaged across *R-1* blocks, having left out block *r*. *M* is the 1984 × 993 design matrix, with each column containing an intercept column as well as a single column for every connection, containing 1 for ME and −1 for idle for the corresponding connection and zeros for the rest of the connections. *β_r_* and *σ_r_* are the 992 × 1 vectors of regression coefficients and errors for the r^th^ fold of the cross-validation respectively. The average contrast pattern was acquired by averaging the regression coefficients over all the cross-validation folds; 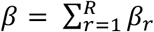. In the resulting average β vector, each regression coefficient *β_l_* (*l* ∈ 1:992) would represent the cross-validated difference or the contrast between the means of its corresponding connections between the two conditions. The links of the *NoI* were considered the 30 highest ranked *c_ij_* (*i* ≠ *j*) in terms of the absolute value of the corresponding *β_l_*.

Moreover, since the absolute value of the contrast is considered to generate the *NoI*, the resulting network reflects changes both in terms of integration (strengthening of a link) and/or segregation (weakening of a link) from one state to the other. The same calculation was carried out in all frequency bins of size 3 to 10 Hz, in the range of 7-40Hz. The optimal frequency bin was considered one that maximized mean absolute contrast over the whole *NoI* network in that band, i.e. 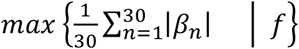, where *f* represents the frequency band in question.

The data collected during the screening session was also used to generate the MI-ME RSN. This network was generated in the exact same cross-validated fashion as the ME-idle RDN i.e. the *NoI*. Only this time around we are interested in the similarity between the two states of ME. Therefore, the MI-ME RSN was generated by sorting the connections in terms of their corresponding *βs* in ascending order. By the same token, the frequency band of interest was considered 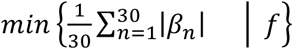.

The MI-ME net similarity in the MI-ME RSN was utilized as a screening tool to monitor the session-to-session effect of training in shifting MI towards ME. MI-ME net similarity was defined as shown in (11)

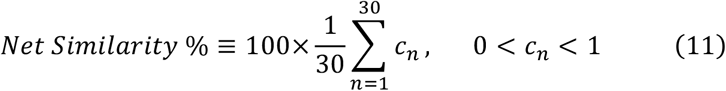

where *c_n_* represents *n*th the link in the RSN.

#### 2.5.2. BCI Training sessions

Each BCI training session took approximately 75 minutes and consisted of two phases; calibration and training, and post training MI and ME cued blocks.

##### Calibration

###### Sub-network library construction

This step of the calibration process is performed off-session to shorten calibration. Based on our pilot experiments, depending on the “*similarity level*” of MI and ME, controlling the 30 links of the *NoI* (i.e. the ME-idle RDN) can be difficult. Therefore, we introduced the concept of “*incremental levels of difficulty*” to the training paradigm. This entailed breaking the 30-link-*NoI* into a number of smaller sub-networks (sub-*NoI*s), and gradually increasing the difficulty, by expanding the training network one sub-*NoI* at a time. The number of links in the sub-*NoI*s was set to six through our pilot experiments, i.e. five 6-link sub-*NoI*s (Figure 3, b-f).

**Figure 3:**
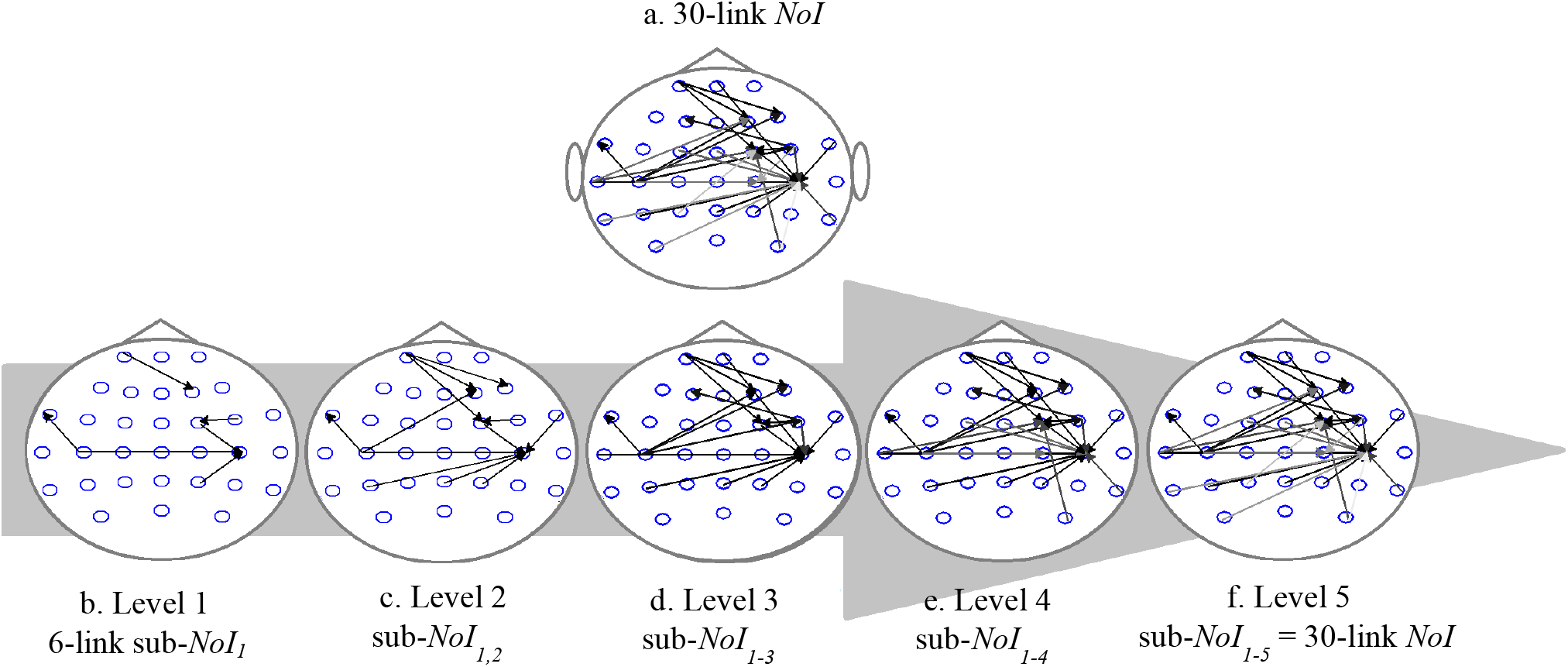
(a.) 30-link ME-idle representational dissimilarity network (RDN), i.e. the *NoI* for subject S1 on BCI session1. b. to f. are levels 1 to 5 of difficulty for controlling the BCI respectively. With each level of progression, one 6-link sub-*NoI* is added to the training network, building up to the 30-link *NoI.*

Identifying the optimal combination of sub-*NoI*s entailed generating all possible 6-link sub-*NoI*s and ranking them in terms of MI-detection performance. Considering the enormous number of sub-*NoI*s (given by the binomial coefficient 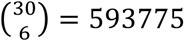), the on-session calibration would become excessively long. To avoid this, a library, narrowed down to a subset of 50 sub-*NoI*s, was constructed at the end of each session and utilized for calibration during the subsequent session. To this end, a 3-minute block consisting of alternating MI-idle trial as well as a 3-minute block of alternating ME-idle trials were recorded at the end of each session (Figure 1).

At the beginning of each BCI training session, the *NoI* was updated using the newly recorded ME data. An individual Mahalanobis minimum distance (MD) classifier was then trained per each possible 6-link sub-*NoI*. The training features for the classifiers consisted of the *DTF* values associated with the 6 links of the corresponding sub-*NoI*, calculated from 1-sec long windows of data while performing ME. Each classifier was then applied to the data from the MI trials for imagery detection. Subsequently, the *NoI*s were ranked based on their accuracies. The top 50 sub-*NoI*s with the highest ranks were selected as members of the sub-*NoI* library, contingent upon every link of the *NoI* being present in at least 8 sub-*NoI*s. This library was then utilized for calibration at the beginning of the next session.

The 5-minute calibration, at the beginning of each BCI session, consisted of a 3-minute block of alternating cued MI-idle trials and a 2-minute classifier update. The on-session recorded MI data, coupled with the sub-*NoI* library from the previous session were utilized to determine the optimal subset of sub-*NoI*s for each session's training. To this end, the 50 sub-*NoI*s in the library were first ranked based on the MI-detection accuracy, using an MD classifier per network. Then, five sub-*NoI*s with the highest MI-detection accuracy that also satisfied inclusion of all the links of the *NoI*, were considered the winning sub-*NoI*s. These networks were then recruited for training with increasing levels of difficulty.

##### Training

Each participant performed six 5-minute blocks of the BCI game, interleaved with a minimum of 2-minute rest blocks. Training on the first BCI session started with the “*easiest*” sub-*NoI*, i.e. the sub-*NoI* ranked with the highest accuracy. The difficulty of controlling the interface would increase by adding one sub-*NoI* at a time gradually building up to the “*hardest*” level i.e. the 30-link *NoI*. The criteria to progress to the next level was the subject-perceived accuracy of 75% on self-report and the actual accuracy of 80%. This approach ensured that the subject perceived sufficient control over the system at each level and felt comfort (Figure 1).

At the beginning of each training session, the level of difficulty was set at one below the last level the subject successfully completed in the previous session. If a subject did not progress during a session, training in the subsequent session started from the easiest level. The maximum number of BCI training sessions was set to 7 *a priori.* However, the training could end with less sessions upon achieving an accuracy of 75% on self-report and an actual accuracy of 80% on the “*hardest*” level (30-link *NoI*).

The VR interface, developed in-house using VRML, consisted of a rubber ball attached to a pipe. A balloon was placed on the tip of the pipe. The subject was asked to imagine repeatedly squeezing on the rubber ball with their non-dominant hand to blow air into the balloon. Every time an MI was detected, the balloon would get bigger. Once a balloon was filled, it would detach from the pipe and be replaced with a balloon of a different color (Figure 4). The MI performance during BCI training was self-paced meaning that the participants performed MI in arbitrary intervals, interleaved with idle. Participants were instructed to keep MI intervals between 4-10 seconds, and to say “*GO*” and “*STOP*” upon starting and stopping imagining respectively.

**Figure 4:**
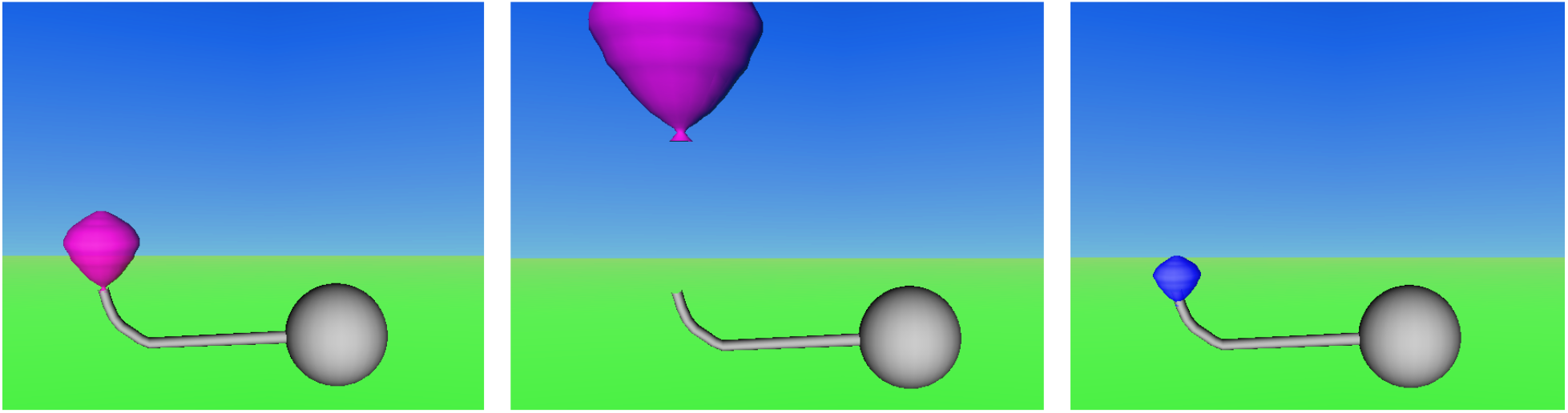
The BCI interface. The subject had to imagine repeatedly squeezing the rubber ball (grey) to blow air into the balloon. Every time an MI was detected, the balloon would expand (left). Once a balloon was filled, it would detach from the pipe (middle) and would be replaced by a balloon of a different color (right).

##### Network update trials

A 3-minute block of alternating MI-idle trials as well as a 3-minute block of alternating ME-idle trials were recorded at the end of each session. These trials were used to update the *NoI* and construct the sub-*NoI* library (see *Calibration*). Moreover, the same data were used to update the MI-ME RSN, used for screening the effect of training, throughout the paradigm. The new ME data were not recorded at the beginning of the session in order to rule out the possibility of the subject having fresh kinesthetic memory of the movement.

#### 2.5.3. Retention session

For every participant, a retention session was scheduled between 10-12 days after their last training session. The data from the ME trials, recorded during the final training session were used for calibration (see *Calibration*). The participant was asked to control the interface starting from the highest level achieved during training. If the accuracy was equal to or more than 75% on self-report and 80% actual accuracy, the changes were considered fully retained. If the subject did not demonstrate this level of accuracy, they were asked to control the interface at consecutively lower levels until either the criteria were met or there were no additional levels. At the end of the retention session, two 7-minute blocks of cued screening trials (alternating MI-idle and alternating ME-idle trials) were recorded.

## 3. Results

Table 1 shows a summary of outcome measures for all subjects. The optimal frequency band for *NoI* generation was within the beta range almost for all subjects (group mean of 16.66 ± 3.51 Hz). Figure 5 shows the MI-ME net similarity from screening session, throughout training and on the retention session. As expected, the results show a considerable inter-subject variability in MI-ME net similarity before and after training (group mean of 32.55% ± 22.90% on the screening session and 55.92% ± 20.90% on the retention session).

**Table 1:**
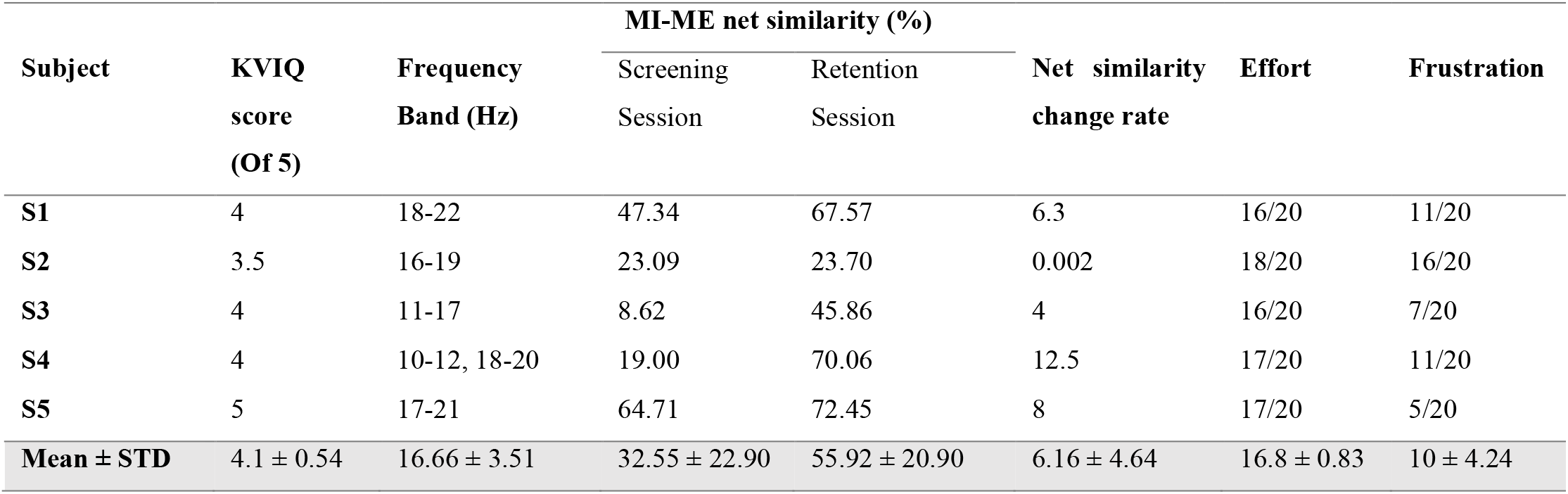
Summary of outcome measures

| Subject | KVIQ<br>score<br>(Of 5) | Frequency<br>Band (Hz) | MI-ME net similarity (%) |  | Net similarity<br>change rate | Effort | Frustration |
| --- | --- | --- | --- | --- | --- | --- | --- |
|  |  |  | Screening<br>Session | Retention<br>Session |  |  |  |
| S1 | 4 | 18-22 | 47.34 | 67.57 | 6.3 | 16/20 | 11/20 |
| S2 | 3.5 | 16-19 | 23.09 | 23.70 | 0.002 | 18/20 | 16/20 |
| S3 | 4 | 11-17 | 8.62 | 45.86 | 4 | 16/20 | 7/20 |
| S4 | 4 | 10-12, 18-20 | 19.00 | 70.06 | 12.5 | 17/20 | 11/20 |
| S5 | 5 | 17-21 | 64.71 | 72.45 | 8 | 17/20 | 5/20 |
| Mean ± STD | 4.1 ± 0.54 | 16.66 ± 3.51 | 32.55 ± 22.90 | 55.92 ± 20.90 | 6.16 ± 4.64 | 16.8 ± 0.83 | 10 ± 4.24 |

**Figure 5:**
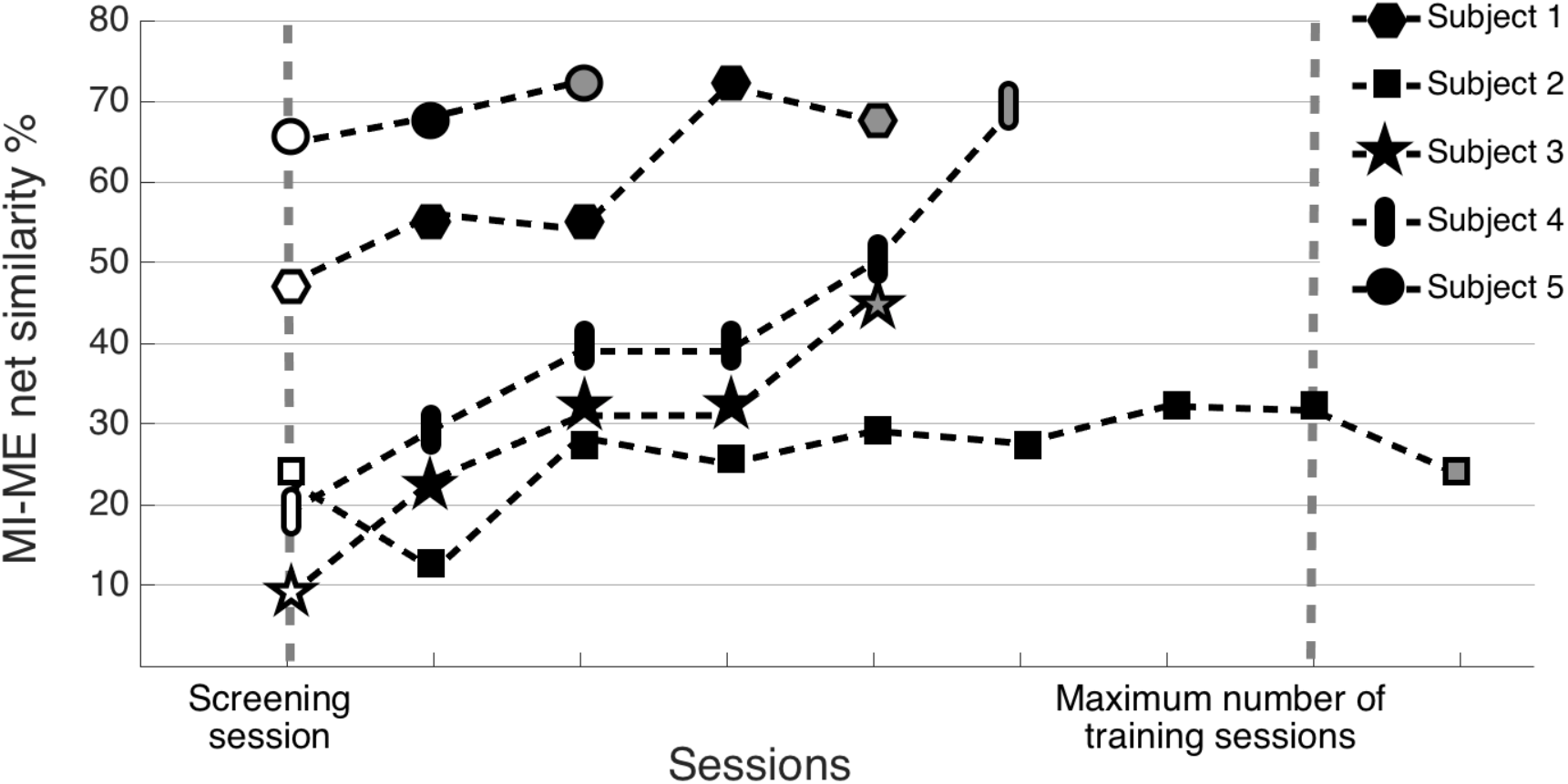
MI-ME net similarity calculated from the MI-ME representational similarity network (RSN) in the absence of feedback. The first (unfilled) and last (grey filled) sessions are screening and retention sessions respectively. The middle sessions (black) are the BCI training sessions.

This inter-subject variability is also present with regards to changing rate of MI-ME net similarity as a result of training (group mean 6.16 ± 4.46). However, except for one, subjects were able to drive the BCI showing a session-to-session increase in their MI-ME net similarity (Figure 5), met the training stopping criteria before 7 sessions (group mean of 3.6 ± 2.1 sessions) and passed the full-retention criteria on the retention session, i.e. controlling the VR environment with on or above 80% accuracy on the “*hardest*” level. Out of 5 recruited participants, S2 (oldest participant: female, aged 56 years) made no progress in operating the BCI system and interestingly, was the only participant who showed no meaningful change in the MI-ME net similarity (Net similarity change rate of 0.002). Subject 5 was the only subject who was able to operate the BCI system with an accuracy of 85% on the first BCI training session, and therefore required no further training.

Figure 6 shows the session-by-session changing process of the MI-Idle RDN towards engaging ME related networks to a higher extent for S1. Figure 6 (a.) shows the ME-idle RDN averaged across all sessions. Figure 6 (b., c., d., e. and f.) show MI-idle RDN on the screening session, first, second and third BCI training sessions as well as the retention session respectively. The shift of MI-idle RDN towards ME-idle RDN is validated by the enhancement in the MI-ME net similarity of 47.34%, 56.01%, 54.12%, 71.81% and 67.57% for Figure 6 (b, c, d, e and f) respectively.

**Figure 6:**
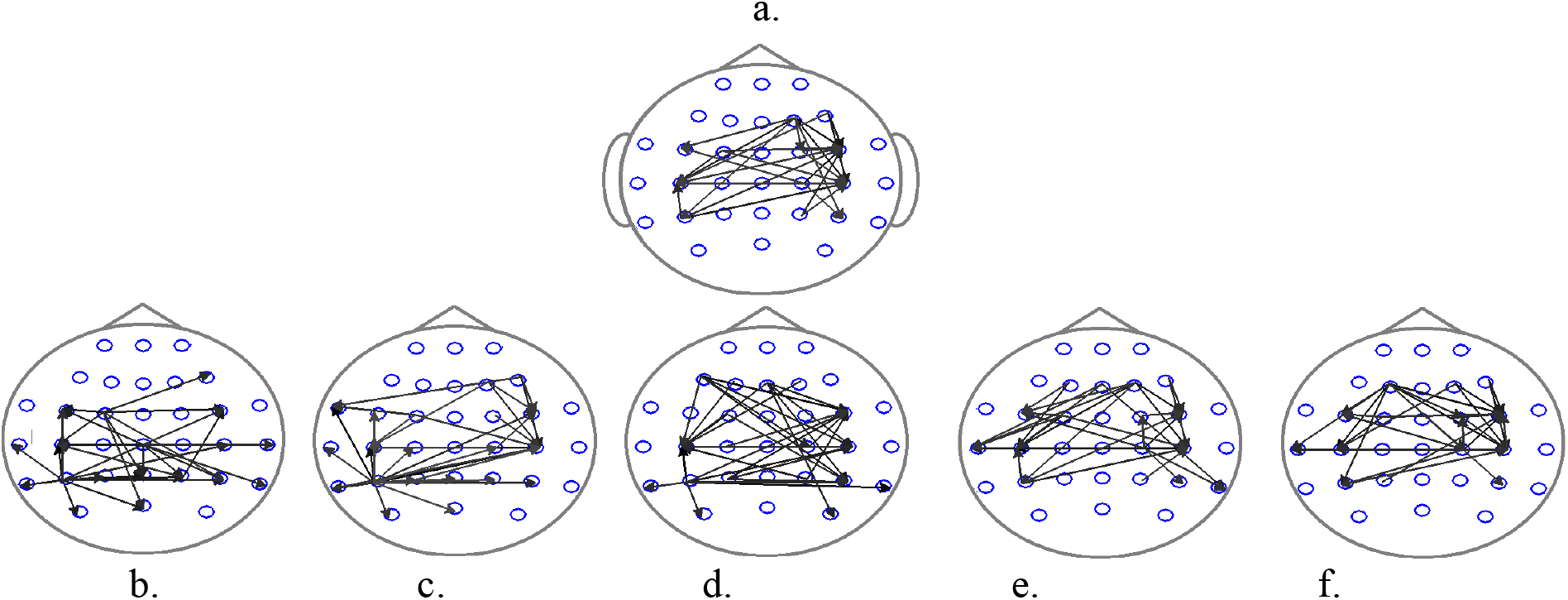
From participant S1: a. Movement execution (ME)-idle Representational dissimilarity network (RDN) averaged across all sessions. b-f. Movement imagination (MI)-idle RDN on the screening session (b), first (c), second (d) and third (e) BCI training sessions as well as the retention (f) session, with MIME net similarity of 47.34%, 56.01%, 54.12%, 71.81% and 67.57% respectively.

## 4. Discussion

Intense practice is required to re-learn the impaired or lost motor skills following a stroke or other brain injury [53]. MI is a mental motor practice that activates the sensorimotor related networks [54] and therefore can serve as an alternative training pathway for patients with higher levels of disability [55]. The first studies investigating the effectiveness of such approaches in post-stroke motor recovery are promising [22, 56–59]. However, the significance of functional changes specific to these interventions is still a matter of debate.

Moreover, when compared to robot-assisted physical therapy in matching doses, BCI interventions fall short in terms of producing functional outcome [22]. There are two main arguments keeping BCI from progressing beyond its current limits as a motor rehabilitation tool that shape the motivation of this study. First, although MI and ME have been shown to engage partially similar neural networks [60], there are fundamental representational dissimilarities between the two states, especially in the primary motor areas [61].

The second argument is the high amount of intra-subject variability in the MI quality [62]. When one is instructed to overtly execute a movement, the quality and consistency of the movement e.g. kinetics, kinematics, can be closely monitored and measured throughout the trials, and modifying instructions can be given as needed. This in turn results in a focused and targeted pattern of activity in the brain as well as the engaged muscle groups, making the outcome of training predictable. MI however, is a purely mental task and therefore highly subjective, making the control for task “correctness” and consistency challenging. This leads to discrepancy in the task-related brain activity across trials [63]. This constrains the functional outcome of BCI by affecting the robustness of the training.

The aim of this study therefore, was to bridge the gap between the network-level representations of movement imagination and execution. This entails “*shifting*” imagination quality in such a way that performing MI engages ME-related networks to a higher extent. In conventional BCIs, i.e. spectral band power-based, the self-modulation of sensorimotor oscillations during MI is supported by providing a type of real-time feedback, in response to the user's alteration of mental state [64]. In this study however, a successful MI was re-defined as an MI that results in an ME-like representation in the brain on from a network perspective. To incorporate this new definition in the BCI design, we propose a connectivity-based real-time MI detection paradigm. The proposed paradigm utilizes the ME-idle representational dissimilarity network (RDN) as a map of reference for MI. Meaning that during training, the brain activity is constantly compared to the ME-idle RDN i.e. the network of interest (*NoI*). Feedback is returned to the subject only when MI elicits a similar pattern of activity in the brain.

Another novelty of the proposed paradigm, was incorporating the idea of “*levels of difficulty*” into the design of the BCI [65]. Training started with a 6-link sub-*NoI*, and upon progression to higher levels, the network was built up, one sub-network at a time, to the 30-link *NoI*. The idea originated from our pilot studies where subjects struggled to drive the 30-link NoI to start with. This design has the advantage of lowering subject frustration, as the mental demands of the task get distributed between 5 levels. The difficulty increased only when the subject felt comfortable with the current level of challenge. Additionally, this design adds an element of flow to the game, which in turn adds to the subject's motivation and engagement [66, 67].

Our results show that four out of five subjects showed enhancement in MI-ME net similarity on a session to session basis as a result of training (Figure 5, Figure 6). Note that the MI-ME representational similarity network (RSN) was updated every session using the cued trials recorded at the end of the session i.e. the network update trials (Figure 1).

Moreover except for S2, subjects were able to reach the training stopping criteria, i.e. on or above 75% self-reported and 80% actual accuracy on the “*hardest*” level. This means that subjects were able to reproduce the ME-idle RDN i.e. the *NoI*, through imagination, at least 80 percent of the time.

The BCI utilized Mahalanobis minimum distance classifier to detect MI, which is the same metric used to generate the *NoI*, as well as the MI-idle RDN and subsequently MI-ME RSN. Therefore, the classification accuracy can be considered a direct reflection of MI-ME net similarity in real-time (see (11)). This would mean that the net similarity in the offline MI-ME RSN after training is expected be close to 80%.

Interestingly, all the subjects failed to reproduce the same level of MI-ME similarity, in the offline trials, with subject-dependent levels of deterioration (Figure 5). This can be interpreted as follows: the enhancement seen in the MI-ME net similarity potentially originates from two components; 1- a reward-dependent or short-term component, which is contingent on receiving the BCI feedback, and 2- a learning or long-term component, as a mental shift in the very way the participant imagines movement. The later component on MI-ME similarity enhancement is retained independent of the feedback, which is depicted in Figure 5. The results suggest that with progression of training with the proposed paradigm, the reward dependent component translates into long-lasting qualitative changes in MI in successful subjects.

This finding is further confirmed in the case of subject S2, i.e. the only subject not able to operate the BCI system (Figure 5, Table 1). S2 was the oldest participant (56 years of age), had the lowest KVIQ score and showed no consistency in the ability to control the virtual reality environment. Intriguingly, S2 was also the only participant who showed no session-to-session enhancement in MI-ME net similarity. This provides evidence that the enhanced similarity seen between imagination and execution is indeed an outcome of training using the proposed paradigm, and not merely a result of MI repetition. Furthermore, this confirms the importance of feedback accuracy in the outcome of BCI training. This is despite the fact that in the offline analysis of the S2's data, modulation of sensorimotor rhythm is detectable as a result of MI. This further validates the argument that even given that the participant is able to control a BP-based MI-BCI, does not ensure a functional outcome as a result of training.

This is in line with the studies investigating the superiority of the general outcome when receiving feedback in response to MI in a BCI paradigm. This is important, since it is known that repetition of a task is sufficient to drive changes in brain activity. Therefore, MI as a stand-alone task, is being incorporated in exercise routines for stroke rehabilitation [68]. As such, some studies have tried to tease apart the BCI-induced changes from the effect of MI repetition. It has been shown that compared to MI alone, neurofeedback-guided MI results in enhanced laterality [69] and magnitude [70] in the sensorimotor rhythms in healthy subjects.

Moreover, in stroke-affected populations, higher functional improvements have been reported in the MIBCI group compared to the control group that received MI training without [59]. Similar outcomes have been reported when the control group received random i.e. sham feedback in response to MI [70]. These studies demonstrated that BCI, if employed prior to physiotherapy, might stimulate brain plasticity. In that sense, BCI can act as a priming mechanism to increase the response of neural networks to physical therapy and thus improve the motor outcome in general [58, 59, 71]. However, neither provide evidence regarding the behavioral specificity of BCI training, reporting a benefit in combined hand and arm [58] or general upper limb [59] Fugl-Meyer assessment score. Additionally, they report no improvement in the subjects’ ability to control the BCI over 3 to 4 weeks of training.

These findings provide further evidence that MI of a specific movement (e.g. opening and closing of the hand) can trigger broad and dispersed neural activity in motor-related areas. This non-specificity in neural activity, limits the improvement in both the functional outcome and BCI performance from a Hebbian standpoint. Furthermore, the large inter-subject variability in the MI-ME net similarity at both screening and retention sessions (Figure 5, Table 1) point to the same aforementioned fact. This is despite the fact that all participants received the exact same instructions for MI performance and they scored the vividness of their kinesthetic MI fairly close to each other. These findings point to the critical need for a “*gold standard*” reference frame, as a measure for MI quality monitoring, apart from self-report. To our knowledge, this is the first study taking a systematic approach to address the aforementioned issue, proposing a priming mechanism to be utilized pre-BCI training.

Moreover, the proposed paradigm offers low computational workload, robust accuracy and is completely self-paced, meaning participants are given the full benefit of using their own intent. Our results provide evidence that the proposed paradigm can effectively induce consistency in MI quality and thus enhance exercise repeatability. This would in turn, result in a more robust training routine. This could potentially help reduce the inter-subject heterogeneity in BCI performance and outcome, and lend to the clinical acceptability of BCI as a neurorehabilitation tool. This makes the outlook of producing a significant functional outcome through BCI training more feasible.

However, a randomized controlled study has yet to be established to investigate the effect of conventional MI-BCI training (sensorimotor rhythm-based BCI) with and without using the proposed priming mechanism. Additionally, it should be noted that this study was conducted with healthy participants who are able to execute overt movements, rendering the ME-related network readily accessible through EEG. However, the target population for the proposed MI-BCI would be patients left with little or no movement residual. This would raise the question of how to generate the ME-idle RDN that was utilized in the proposed paradigm to drive the VR interface from such clinical populations. It has been shown that passive movement accompanied with movement imagination best matches overt movement execution in terms of neural activity in both healthy and stroke subjects [72]. In continuation of this study therefore, we will investigate the plausibility of using passive movement + MI as a substitute for ME in healthy and stroke participants. Furthermore, the ability of stroke affected participants to control sensory motor rhythm BCI via movement imagination has been demonstrated in multiple studies [2, 22, 73, 74]. However, the ability of stroke patients to operate the proposed network-based BCI that defines a successful MI, one that engages ME-related networks, needs further investigation.

## Acknowledgment

We thank the participants who took part in this study.

## Reference

[1] Pfurtscheller G, Müller-Putz G, Scherer R. Rehabilitation with brain-computer interface systems. Computer (Long Beach Calif) http://ieeexplore.ieee.org/abstract/document/4640664/ (2008, accessed 6 April 2017).

[2] Nilsen D, Gillen G, Gordon A. Use of mental practice to improve upper-limb recovery after stroke: a systematic review. Am J Occup http://ajot.aota.org/Article.aspx?articleid=1854525 (2010, accessed 6 April 2017).

[3] Jeannerod M, Decety J. Mental motor imagery: a window into the representational stages of action. Curr Opin Neurobiol http://www.sciencedirect.com/science/article/pii/0959438895800999 (1995, accessed 7 April 2017).

[4] Decety J. Do imagined and executed actions share the same neural substrate? Cogn brain Res http://www.sciencedirect.com/science/article/pii/092664109500033X (1996, accessed 7 April 2017).

[5] Ehrsson H, Geyer S, Naito E. Imagery of voluntary movement of fingers, toes, and tongue activates corresponding body-part-specific motor representations. J Neurophysiol http://jn.physiology.org/content/90/5/3304.short (2003, accessed 7 April 2017).

[6] Gao Q, Duan X, Chen H. Evaluation of effective connectivity of motor areas during motor imagery and execution using conditional Granger causality. Neuroimage http://www.sciencedirect.com/science/article/pii/S1053811910011687 (2011, accessed 7 April 2017).

[7] Scott S. Optimal feedback control and the neural basis of volitional motor control. Nat Rev Neurosci http://www.nature.com/nrn/journal/v5/n7/abs/nrn1427.html (2004, accessed 7 April 2017).

[8] Brontë-Stewart H, Lisberger S. Physiological properties of vestibular primary afferents that mediate motor learning and normal performance of the vestibulo-ocular reflex in monkeys. J Neurosci http://www.jneurosci.org/content/14/3/1290.short (1994, accessed 7 April 2017).

[9] Dimitriou M. Enhanced Muscle Afferent Signals during Motor Learning in Humans. Curr Biol http://www.sciencedirect.com/science/article/pii/S0960982216300835 (2016, accessed 7 April 2017).

[10] Silvoni S, Ramos-Murguialday A, Cavinato M, et al. Brain-computer interface in stroke: A review of progress. Clin EEG Neurosci; 42: 245–252.

[11] Gomez-Rodriguez M, Peters J, Hill J, et al. Closing the sensorimotor loop: Haptic feedback facilitates decoding of arm movement imagery. In: Systems Man and Cybernetics (SMC), 2010 IEEE International Conference on. IEEE, 2012, pp. 121–126.

[12] Sharma N, Pomeroy VM, Baron J-C. Motor Imagery A Backdoor to the Motor System After Stroke? Stroke 2006; 37: 1941–1952.

[13] Langhorne P, Coupar F, Pollock A. Motor recovery after stroke: a systematic review. Lancet Neurol http://www.sciencedirect.com/science/article/pii/S1474442209701504 (2009, accessed 8 April 2017).

[14] Ietswaart M, Johnston M, Dijkerman H, et al. Mental practice with motor imagery in stroke recovery: randomized controlled trial of efficacy. Brain http://brain.oxfordjournals.org/content/early/2011/04/22/brain.awr077.short (2011, accessed 8 April 2017).

[15] Creem-Regehr S. Sensory-motor and cognitive functions of the human posterior parietal cortex involved in manual actions. Neurobiol Learn Mem http://www.sciencedirect.com/science/article/pii/S1074742708001883 (2009, accessed 8 April 2017).

[16] Haller S, Chapuis D, Gassert R. Supplementary motor area and anterior intraparietal area integrate fine-graded timing and force control during precision grip. Eur J http://onlinelibrary.wiley.com/doi/10.1111/j.1460-9568.2009.07003.x/full (2009, accessed 8 April 2017).

[17] Bauer R, Fels M, Vukelić M, et al. Bridging the gap between motor imagery and motor execution with a brain–robot interface. Neuroimage http://www.sciencedirect.com/science/article/pii/S1053811914010180 (2015, accessed 8 April 2017).

[18] Johnson-Frey S, Newman-Norlund R. A distributed left hemisphere network active during planning of everyday tool use skills. Cereb cortex http://cercor.oxfordjournals.org/content/15/6/681.short (2005, accessed 8 April 2017).

[19] Grefkes C, Eickhoff S, Nowak D, et al. Dynamic intra-and interhemispheric interactions during unilateral and bilateral hand movements assessed with fMRI and DCM. Neuroimage http://www.sciencedirect.com/science/article/pii/S1053811908002838 (2008, accessed 8 April 2017).

[20] Shibasaki H. Cortical activities associated with voluntary movements and involuntary movements. Clin Neurophysiol http://www.sciencedirect.com/science/article/pii/S1388245711005463 (2012, accessed 8 April 2017).

[21] Chouinard P, Paus T. The primary motor and premotor areas of the human cerebral cortex. Neurosci http://journals.sagepub.com/doi/abs/10.1177/1073858405284255 (2006, accessed 8 April 2017).

[22] Ang KK, Guan C, Chua KSG, et al. A large clinical study on the ability of stroke patients to use an EEG-based motor imagery brain-computer interface. Clin EEG Neurosci 2011; 42: 253–258.

[23] Kaiser V, Kreilinger A, Müller-Putz G. First steps toward a motor imagery based stroke BCI: new strategy to set up a classifier. Front http://journal.frontiersin.org/article/10.3389/fnins.2011.00086 (2011, accessed 8 April 2017).

[24] Carter AR, Shulman GL, Corbetta M. Why use a connectivity-based approach to study stroke and recovery of function? Neuroimage 2012; 62: 2271–2280.

[25] Silasi G, Murphy T. Stroke and the connectome: how connectivity guides therapeutic intervention. Neuron http://www.sciencedirect.com/science/article/pii/S089662731400782X (2014, accessed 8 April 2017).

[26] Doyon J, Benali H. Reorganization and plasticity in the adult brain during learning of motor skills. Curr Opin Neurobiol http://www.sciencedirect.com/science/article/pii/S095943880500036X (2005, accessed 8 April 2017).

[27] Fornito A, Zalesky A, Breakspear M. The connectomics of brain disorders. Nat Rev Neurosci http://www.nature.com/nrn/journal/v16/n3/abs/nrn3901.html (2015, accessed 8 April 2017).

[28] Mawase F, Uehara S, Bastian A. Motor learning enhances use-dependent plasticity. J http://www.jneurosci.org/content/37/10/2673.abstract (2017, accessed 8 April 2017).

[29] Mokienko O, Chernikova L. Motor imagery and its practical application. Neurosci http://search.proquest.com/openview/72cfb214d748ad73e34d43b8a3d7c49d/1?pqorigsite=gscholar&cbl=38002 (2014, accessed 9 April 2017).

[30] Holpera L, Kobashib N, Kiperb D, et al. Trial-to-trial variability differentiates motor imagery during observation between low versus high responders: A functional near-infrared spectroscopy study. Behav Brain Res 2012; 229: 29–40.

[31] Miller L, Saygin A. Individual differences in the perception of biological motion: links to social cognition and motor imagery. Cognition http://www.sciencedirect.com/science/article/pii/S0010027713000656 (2013, accessed 9 April 2017).

[32] Holz E, Höhne J, Staiger-Sälzer P. Brain–computer interface controlled gaming: Evaluation of usability by severely motor restricted end-users. Artif Intell http://www.sciencedirect.com/science/article/pii/S0933365713001140 (2013, accessed 9 April 2017).

[33] Arvaneh M, Guan C, Ang KK, et al. Robust EEG channel selection across sessions in brain-computer interface involving stroke patients. In: Proceedings of the International Joint Conference on Neural Networks. Brisbane, QLDhttp://www.scopus.com/inward/record.url?eid=2-s2.0-84865090992&partnerID=40&md5=5ea3a530a3e2284b0b14e0858c875905.

[34] Friston K. Functional and effective connectivity: a review. Brain Connect http://online.liebertpub.com/doi/abs/10.1089/brain.2011.0008 (2011, accessed 9 April 2017).

[35] Honey C, Sporns O, Cammoun L. Predicting human resting-state functional connectivity from structural connectivity. Proc http://www.pnas.org/content/106/6/2035.short (2009, accessed 9 April 2017).

[36] Schoffelen J, Gross J. Source connectivity analysis with MEG and EEG. Hum Brain Mapp http://onlinelibrary.wiley.com/doi/10.1002/hbm.20745/full (2009, accessed 9 April 2017).

[37] Sakkalis V. Review of advanced techniques for the estimation of brain connectivity measured with EEG/MEG. Comput Biol Med 2011; 41: 1110–1117.

[38] Daly I, Nasuto S, Warwick K. Brain computer interface control via functional connectivity dynamics. Pattern Recognit http://www.sciencedirect.com/science/article/pii/S0031320311002032 (2012, accessed 9 April 2017).

[39] Liu M, Kuo C, Chiu A. Statistical threshold for nonlinear granger causality in motor intention analysis. Eng Med http://ieeexplore.ieee.org/abstract/document/6091247/ (2011, accessed 9 April 2017).

[40] Billinger M, Brunner C, Scherer R, et al. Towards a framework based on single trial connectivity for enhancing knowledge discovery in BCI. In: Active Media Technology. Springer, 2012, pp. 658–667.

[41] Heger D, Terziyska E, Schultz T. Connectivity based feature-level filtering for single-trial eeg bcis. Acoust Speech Signal http://ieeexplore.ieee.org/abstract/document/6853962/ (2014, accessed 9 April 2017).

[42] Pfurtscheller G, Lopes da Silva FH. Event-related EEG/MEG synchronization and desynchronization: basic principles. Clin Neurophysiol 1999; 110: 1842–1857.

[43] Cadotte AJ, DeMarse TB, He P, et al. Causal measures of structure and plasticity in simulated and living neural networks. PLoS One 2008; 3: e3355.

[44] Chen H, Yang Q, Liao W, et al. Evaluation of the effective connectivity of supplementary motor areas during motor imagery using Granger causality mapping. Neuroimage 2009; 47: 1844–1853.

[45] Sameshima K, Baccalá L. Using partial directed coherence to describe neuronal ensemble interactions. J Neurosci Methods http://www.sciencedirect.com/science/article/pii/S0165027099001284 (1999, accessed 9 April 2017).

[46] Vicente R, Wibral M, Lindner M, et al. Transfer entropy a model-free measure of effective connectivity for the neurosciences. J Comput Neurosci 2011; 30: 45–67.

[47] Kaminski MJ, Blinowska KJ. A new method of the description of the information flow in the brain structures. Biol Cybern 1991; 65: 203–210.

[48] Astolfi L, Cincotti F, Mattia D, et al. Comparison of different cortical connectivity estimators for high resolution EEG recordings. Hum Brain Mapp 2007; 28: 143–157.

[49] Babiloni F, Cincotti F, Babiloni C, et al. Estimation of the cortical functional connectivity with the multimodal integration of high-resolution EEG and fMRI data by directed transfer function. Neuroimage http://www.sciencedirect.com/science/article/pii/S1053811904005646 (2005, accessed 9 April 2017).

[50] Wang J, Zuo X, Gohel S, et al. Graph theoretical analysis of functional brain networks: test-retest evaluation on short-and long-term resting-state functional MRI data. PLoS One http://journals.plos.org/plosone/article?id=10.1371/journal.pone.0021976 (2011, accessed 9 April 2017).

[51] Vinck M, Oostenveld R, Wingerden M Van, et al. An improved index of phase-synchronization for electrophysiological data in the presence of volume-conduction, noise and sample-size bias. Neuroimage http://www.sciencedirect.com/science/article/pii/S1053811911000917 (2011, accessed 9 April 2017).

[52] Hoerl A, Kennard R. Ridge regression: Biased estimation for nonorthogonal problems. Technometrics http://amstat.tandfonline.com/doi/abs/10.1080/00401706.1970.10488634 (1970, accessed 9 April 2017).

[53] Halsband U, Lange R. Motor learning in man: a review of functional and clinical studies. J Physiol http://www.sciencedirect.com/science/article/pii/S0928425706000155 (2006, accessed 9 April 2017).

[54] Hyde C, Fuelscher I, Lum J, et al. Primary Motor Cortex Excitability Is Modulated During the Mental Simulation of Hand Movement. J https://www.cambridge.org/core/journals/journal-of-the-international-neuropsychological-society/article/div-classtitleprimary-motor-cortex-excitability-is-modulated-during-the-mental-simulation-of-hand-movementdiv/EEA789EFD7CB50E55734537AC26F1027 (2017, accessed 7 April 2017).

[55] Szameitat A, Shen S, Conforto A, et al. Cortical activation during executed, imagined, observed, and passive wrist movements in healthy volunteers and stroke patients. Neuroimage http://www.sciencedirect.com/science/article/pii/S1053811912004971 (2012, accessed 9 April 2017).

[56] Prasad G, Herman P, Coyle D, et al. Applying a brain-computer interface to support motor imagery practice in people with stroke for upper limb recovery: a feasibility study. J Neuroeng Rehabil 2010; 7: 60.

[57] Shindo K, Kawashima K, Ushiba J, et al. Effects of neurofeedback training with an electroencephalogram-based brain-computer interface for hand paralysis in patients with chronic stroke: A preliminary case series study. J Rehabil Med 2011; 43: 951–957.

[58] Ramos-Murguialday A, Broetz D, Rea M. Brain–machine interface in chronic stroke rehabilitation: a controlled study. Ann http://onlinelibrary.wiley.com/doi/10.1002/ana.23879/full (2013, accessed 8 April 2017).

[59] Pichiorri F, Morone G, Petti M, et al. Brain–computer interface boosts motor imagery practice during stroke recovery. Ann http://onlinelibrary.wiley.com/doi/10.1002/ana.24390/full (2015, accessed 9 April 2017).

[60] Gerardin E, Sirigu A, Lehéricy S, et al. Partially overlapping neural networks for real and imagined hand movements. Cerebral http://cercor.oxfordjournals.org/content/10/11/1093.short (2000, accessed 9 April 2017).

[61] Higuchi S, Imamizu H, Kawato M. Cerebellar activity evoked by common tool-use execution and imagery tasks: an fMRI study. Cortex http://www.sciencedirect.com/science/article/pii/S001094520870460X (2007, accessed 9 April 2017).

[62] Aliakbaryhosseinabadi S, Kostic V, Pavlovic A. Effect of Attention Variation in Stroke Patients: Analysis of Single Trial Movement-Related Cortical Potentials. Converging Clin http://link.springer.com/chapter/10.1007/978-3-319-46669-9_159 (2017, accessed 20 May 2017).

[63] Jeunet C, N'Kaoua B, Lotte F. Advances in user-training for mental-imagery-based BCI control: Psychological and cognitive factors and their neural correlates. Prog Brain Res http://www.sciencedirect.com/science/article/pii/S0079612316300061 (2016, accessed 9 April 2017).

[64] Vukelić M, Gharabaghi A. Self-regulation of circumscribed brain activity modulates spatially selective and frequency specific connectivity of distributed resting state networks. Front Behav http://journal.frontiersin.org/article/10.3389/fnbeh.2015.00181 (2015, accessed 8 April 2017).

[65] Cameirao MS, Bermudez IBS, Duarte Oller E, et al. The rehabilitation gaming system: a review. Stud Heal Technol Inf 2009; 145: 65–83.

[66] Czikszentmihalyi M. Flow: The psychology of optimal experience. Praha Lidov © Noviny Cited page.

[67] Sweetser P, Wyeth P. GameFlow: a model for evaluating player enjoyment in games. Comput Entertain 2005; 3: 3.

[68] Sharma N, Baron J, Rowe JB. Motor imagery after stroke: relating outcome to motor network connectivity. Ann Neurol 2009; 66: 604–616.

[69] Boe S, Gionfriddo A, Kraeutner S, et al. Laterality of brain activity during motor imagery is modulated by the provision of source level neurofeedback. Neuroimage http://www.sciencedirect.com/science/article/pii/S1053811914005436 (2014, accessed 9 April 2017).

[70] Bai O, Huang D, Fei D, et al. Effect of real-time cortical feedback in motor imagery-based mental practice training. NeuroRehabilitation http://content.iospress.com/articles/neurorehabilitation/nre1039 (2014, accessed 9 April 2017).

[71] Grosse-Wentrup M, Mattia D, Oweiss K. Using brain-computer interfaces to induce neural plasticity and restore function. J Neural Eng 2011; 8: 25004.

[72] Kang H, Park W, Kang J, et al. A neural analysis on motor imagery and passive movement using a haptic device. Syst (ICCAS …http://ieeexplore.ieee.org/abstract/document/6393082/ (2012, accessed 9 April 2017).

[73] Soekadar SR, Birbaumer N, Cohen LG. Brain-computer interfaces in the rehabilitation of stroke and neurotrauma. In: Systems neuroscience and rehabilitation. Springer Tokyo, 2011.

[74] Vries S de, Tepper M, Otten B, et al. Recovery of motor imagery ability in stroke patients. Rehabil Res https://www.hindawi.com/journals/rerp/2011/283840/abs/ (2011, accessed 6 April 2017).

